# Sketching and sampling approaches for fast and accurate long read classification

**DOI:** 10.1101/2021.11.04.467374

**Authors:** Arun Das, Michael C. Schatz

## Abstract

**Background:** In modern sequencing experiments, quickly and accurately identifying the sources of the reads is a crucial need. In metagenomics, where each read comes from one of potentially many members of a community, it can be important to identify the exact species the read is from. In other settings, it is important to distinguish which reads are from the targeted sample and which are from potential contaminants. In both cases, identification of the correct source of a read enables further investigation of relevant reads, while minimizing wasted work. This task is particularly challenging for long reads, which can have a substantial error rate that obscures the origins of each read.

**Results:** Existing tools for the read classification problem are often alignment or index-based, but such methods can have large time and/or space overheads. In this work, we investigate the effectiveness of several sampling and sketching-based approaches for read classification. In these approaches, a chosen sampling or sketching algorithm is used to generate a reduced representation (a “screen”) of potential source genomes for a query readset before reads are streamed in and compared against this screen. Using a query read’s similarity to the elements of the screen, the methods predict the source of the read. Such an approach requires limited pre-processing, stores and works with only a subset of the input data, and is able to perform classification with a high degree of accuracy.

**Conclusions:** The sampling and sketching approaches investigated include uniform sampling, methods based on MinHash and its weighted and order variants, a minimizer-based technique, and a novel clustering-based sketching approach. We demonstrate the effectiveness of these techniques both in identifying the source microbial genomes for reads from a metagenomic long read sequencing experiment, and in distinguishing between long reads from organisms of interest and potential contaminant reads. We then compare these approaches to existing alignment, index and sketching-based tools for read classification, and demonstrate how such a method is a viable alternative for determining the source of query reads. Finally, we present a reference implementation of these approaches at https://github.com/arun96/sketching.

## 1. Background

Metagenomics has become an increasingly popular area of study over the past two decades, and has enabled us to better understand the diversity, interactions and evolution of microbial communities in a plethora of environments (Quince et al. 2017; Handelsman Jo 2004; Florian P. Breitwieser, Lu, and Salzberg 2019). Metagenomics has highlighted the problem of being able to quickly and accurately identify the source of a given DNA sequence from all the genomic material in a given sample. This is needed to classify and sort reads for further downstream analysis, and to identify and remove potential contaminants that are present in a sample. Efficient solutions to such problems are especially important in metagenomics, where the scale of these microbial communities can be extremely large. Individual metagenomics datasets can contain thousands of genomes, and large sequence repositories such as Refseq (Pruitt et al. 2000; W. Li et al. 2021) contain hundreds of thousands of microbial genomes against which metagenomic sequencing reads may need to be compared. The scale of the metagenomics sequencing experiments themselves are also massive; initiatives like the Tara Oceans Project (Sunagawa et al. 2015, 2020),the MetaSUB Research Consortium (Danko et al. 2021) and the Twitchell Wetlands sequencing effort have generated 7.2 trillion, 8 trillion and 2.6 trillion bases of sequencing data respectively across thousands of samples.

The read classification problem is to identify the source genome of a given input read, usually by comparing the read to a list of potential source genomes and choosing the one with the highest similarity. This comparison may be done naively by comparing the entirety of each read to the entirety of each genome to find the best alignment or through an exhaustive analysis of k-mers present. While these approaches are highly accurate they can incur high computational overheads, which presents an opportunity for lower overhead techniques such as sketching or sampling, especially for long read data.

Sketching is the process of generating an approximate, compact summary of the data (a “sketch”), which retains properties of interest and can be used as a proxy for the original data (Rowe 2019).

Sampling selects a subset of the data, either systematically or randomly, but does not guarantee the preservation of these properties. Each has unique advantages: sketching has been shown to bound error better than sampling (Cormode 2017; Rowe 2019), while systematic sampling (such as uniform sampling) can provide bounds on the number of samples from specific sections of the original data included in the generated subset. Both sketching and sampling provide simple routes to greatly reduce the size of an input set, while retaining the characteristics and features that identify the set, thus allowing a comparable level of accuracy.

One of the most well-known sketching approaches, and the main one we employ in our work, is MinHash, which was first presented as a method to estimate document similarity using the similarity between their hashed sub-parts (Broder 1997). It is now widely used in genomics, such as in Mash (Ondov et al. 2016), which performs fast similarity and distance estimation between two input sequences, and tools such as Mash Screen (Ondov et al. 2019) which uses MinHash to predict which organisms are contained in a mixture. Other tools include MashMap (Jain, Dilthey, et al. 2018), which blends minimizers and MinHash for fast, approximate alignment of DNA sequences, and MHAP (Berlin et al.2015) to accelerate genome assembly. Beyond MinHash, several related approaches have been proposed, such as bloom filters (Solomon and Kingsford 2016; Sun et al. 2018), the HyperLogLog sketch (Baker and Langmead 2019; F. P. Breitwieser, Baker, and Salzberg 2018), and other sketching approaches to estimate similarity, containment or cardinality (Marçais, Solomon, et al. 2019).

### 1.1. Approaches to Read Classification

The simplest approach to read classification is to align each query read to all potential source genomes, and using the genome with the best alignment as the predicted source. While the most accurate approach would be exhaustive sequence-to-sequence alignment with dynamic programming, this is impractically slow, so aligners typically use some form of seed-and-extend that start with exact matches and build out longer regions of high similarity. Tools such as Minimap2 (H. Li 2018), Winnowmap (Jain, Rhie, Hansen,et al. 2020) and Winnowmap2 (Jain et al. 2022) use a variation of this approach in the anchor chaining strategy, where sets of exactly matched seeds are chained together to aid in alignment. However, even with these optimizations, alignment still remains computationally expensive, and offers a level of detail not always necessary in read classification.

A more sophisticated approach is index-based analysis, where a pre-computed index is constructed with sequences that are specific or important to each genome or group of interest. Then each query read is classified by the presence or absence of these pre-identified markers. The foremost examples of this form of read classification are the Kraken (Wood and Salzberg 2014; Wood, Lu, and Langmead 2019) set of tools, as well as tools such as CLARK (Ounit et al. 2015) and Centrifuge (Kim et al. 2016). While the read classification process in index-based approaches can be extremely fast, there is substantial time and space overhead associated with the construction of the index.

The space, time and computational overhead associated with alignment- and index-based read classification has motivated the need for faster, more accurate, and lower overhead alternatives. Sketching has proven to be a viable solution instead of whole genome comparisons as it provides the level of accuracy required for less demanding tasks such as read classification, while substantially reducing overhead. Examples of this are MashMap (Jain, Dilthey, et al. 2018) and MetaMaps (Dilthey et al. 2019), which use approximate similarity instead of exact alignment between regions of two sequences to perform alignment.

In this work, we critically evaluate several sketching and sampling methods that aim to reduce the computational overhead of read classification. We apply sketching, using MinHash- and minimizer-based approaches, as well as uniform sampling, to generate compact, approximate representations of potential source genomes for a given readset. We then classify reads against these representations, and demonstrate that we are able to classify, with a high degree of accuracy, reads from a microbial community and detect contaminants in real and simulated sequencing experiments.

## 2. Methods

In our methods, we consider several sketching and sampling approaches to generate reduced representations of the source genomes. We refer to this collection of reduced representations, and any auxiliary information generated alongside them, as a “screen” of the genomes, inspired by the use of the term to describe a collection of MinHash sketches in Mash Screen (Ondov et al. 2019). The screen acts as a proxy for the source genomes, removing the need to store or use the original sequences. In our work, a screen comprises sets of k-mers, one for each potential source genome, with each set of k-mers being the reduced representation for the original genome they were generated from.

Query reads are then compared against this screen, with the read being classified to the most similar reduced representation in the screen **(Figure 1)**. Use of just the screen instead of the genomes themselves significantly reduces the computation necessary to determine the source of a query read.

**Figure 1.**
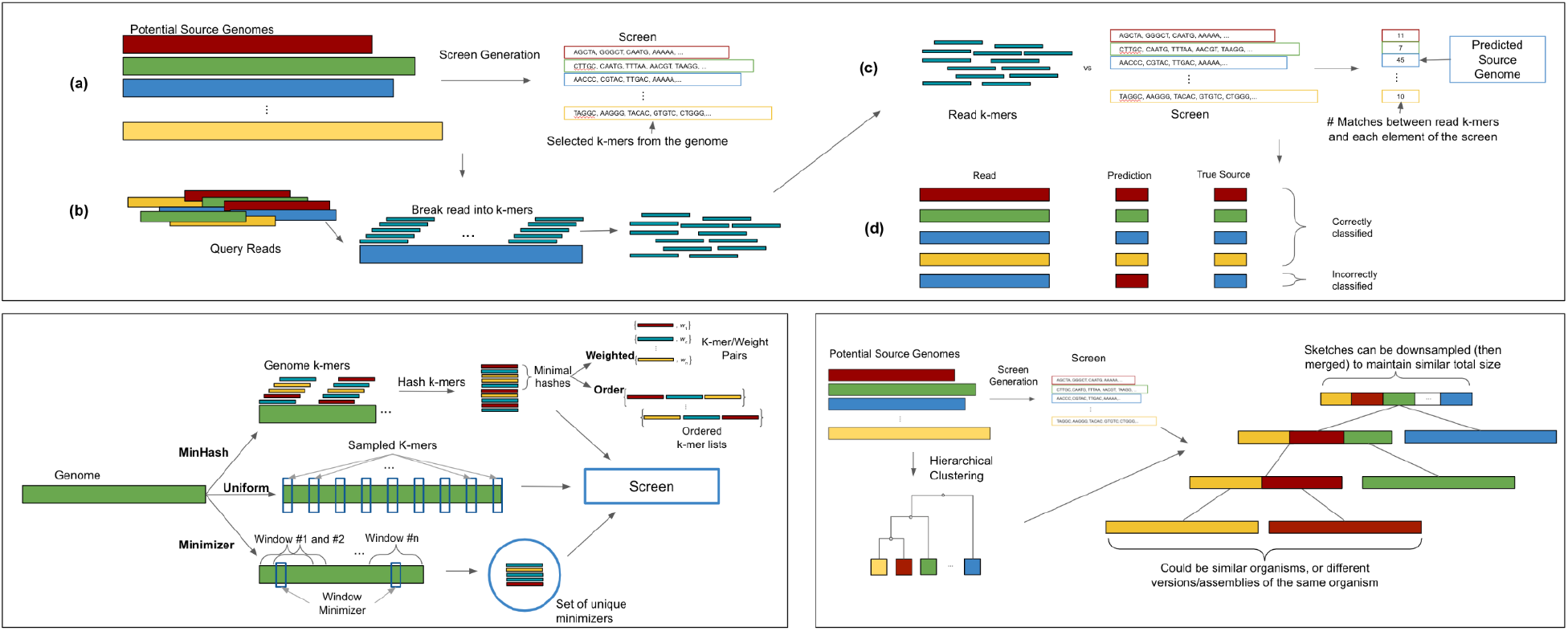
Overview of sketching and sampling methods. **(Top)** The screen is generated using the desired sketching or sampling approach from potential input genomes, read k-mers are compared against the screen, with the element of the screen most similar to the read predicted as its source. **(Bottom Left**) The different sketching and sampling approaches used to generate a screen. **(Bottom Right)** Sketch clustering: input genomes are clustered, and the generated screen is arranged to match this clustering, with reads compared to the root and then down the tree.

### 2.1 Determining sketch and sample size

As our goal is to reduce the computation needed to determine the source of a read, the single biggest factor in such an approach is the size of the screen, or the fraction of the k-mers from the genomes that are stored. The ideal screen size will minimize the input storage requirement while being detailed enough to capture the specificity of each genome. To do this, three main factors must be considered: (1) the size of the genomes (in base pairs); (2) the read length and error rate of the reads we are classifying; and (3) the amount of similarity needed to correctly match a read and its true source genome. We refer to this as the number “target matches” or “shared hashes”, which is the number of sketched or sampled k-mers a read and its source genome share. We formalize this using the following formula:

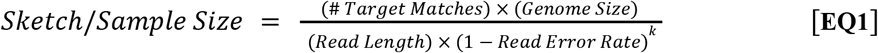

This formula allows us to sketch and sample at a rate where we expect to retain the target number of k-mers per read length of sequence in the original genome, adjusted for error. We adjust for error by computing the fraction of k-mers we expect to be affected by error at that error rate, and oversampling or oversketching to compensate for this. The resulting sketch or sample has, in expectation, the desired number of error-free hashes stored from each read length of sequence in the genome, and therefore the desired number of error-free shared hashes with a read drawn from the same region. This formula is used to determine the expected number of stored hashes in all our approaches, as the cost of generating similar sized sketches and samples is relatively equal across the approaches.

As shown in **EQ1**, the exact size of the screen depends on the experimental parameters. The compression factor is equal to the targeted number of k-mer matches per read divided by the read length, with an oversampling by a factor of *1*/(*1* - *error*)^*k*^ to correct for errors. This makes these approaches best suited for lower error, longer reads, as these require the fewest number of hashes across the genome, with shorter or higher error reads requiring larger screens. The number of target matches also determines screen size, but as we will see in the results section, high accuracy is possible with small screens that are much smaller than the original data, especially for low error rate and longer reads.

During classification, read k-mers are only compared against the stored k-mers in the screen. This can be done efficiently using hash table-based sets: the space required for representing the reference genomes will typically be only a few percent of the total sequence length, and the number of hash table lookups will also be substantially reduced. Once computed, a screen can be re-used for all future runs, amortizing the cost of screen generation across all future uses. This approach also reduces the cost of updating the set of potential source genomes; instead of rebuilding the whole index, sketches or samples can easily be added or removed, with the rest of the screen left unchanged.

### 2.2 Overview of sketching and sampling approaches

In this work, we consider several existing sketching and sampling techniques. Given a list of input genomes, each of these techniques generates a set of selected k-mers to act as a reduced representation of each original sequence. These sets of selected k-mers make up the elements of the screen that all query reads are compared against to determine their source genome. In the case of more sophisticated techniques that generate auxiliary information, such as weights or orderings of the stored k-mers, the screen will also contain this data for use during comparisons of query reads.

In order to allow strand-neutral comparisons, all approaches will use and store canonical k-mers. In our implementation, this is defined as the lexicographically smaller of the forward and reverse complement representations of the k-mer being considered. This is the same definition used in Mash (Ondov et al.2016) and Kraken (Wood and Salzberg 2014).

A reference implementation of these approaches can be found at https://github.com/arun96/sketching.

#### Uniform

In this approach, k-mers are uniformly extracted across the genome to reach the desired sample size. The chief benefit of this approach is simplicity, including a guarantee on the maximum distance between k-mers in our screen, which is generally not guaranteed for alternative approaches. This also guarantees that each read will have a highly predictable amount of overlap with the sampled version of its source genome, though error in these samples can obscure the detection of this overlap. Computationally, uniform sampling is the simplest of the approaches; For a genome of size *n*, a sample of size *s* can be generated by selecting a k-mer every *n/s* bases, meaning the sample can be generated in O(*n*) time and stored in O(*s*) space.

#### MinHash

This sketching technique enables quickly estimating the similarity between two input sequences by computing the Jaccard coefficient of the selected k-mers extracted from one sequence compared to those selected in a second sequence. There are several widely used methods to generate a MinHash sketch, such as using multiple hash functions or a partitioning of the space of possible k-mers. For our analysis, we use a single hash function, and select the *s* smallest hash values returned, such as is used in Mash. As hashing is simply a permutation of the input values, this effectively generates a random sampling of *s* k-mers to be used as the representation of the original genome. In terms of computation, MinHash requires all k-mers to be hashed while maintaining a list of minimal hashes, which can be done using a fixed-size max-heap. A sketch of size *s* from a genome of size *n* requires O(*n*) time to hash each k-mer. Then the *s* smallest hash values can then be identified using a max-heap of size *s* for a total runtime of O(*n* log *s*).

#### Weighted MinHash (WMH)

This enhances a basic MinHash with weights for each k-mer representing a measure of the k-mer’s “importance”, with more highly weighted k-mers indicating a greater level of confidence in a match. Weights are typically based on the number of times that an element occurs, or on a predetermined scoring scheme. In our reference implementation, the weight is a measure of “uniqueness”; we compute the weight of a k-mer as the total number of genomes in our screen minus the number of genomes the k-mer is found in. Unique k-mers occurring in a single genome are weighted the highest as these are strong candidates for precisely identifying the source of a read. This is especially useful when considering reads that share the same number of k-mers with multiple potential genomes; having a shared highly weighted k-mer with one of these potential genomes can help determine the source with more accuracy than a random tie break. This approach can be further extended to add a “multiplier” into weighted MinHash, where unique k-mers have their computed weight multiplied by some multiplier M (M=1 in regular weighted MinHash), allowing these highly informative k-mers to play an even larger role in determining the similarity between a read and its genome.

The addition of weight is computationally expensive; for a sketch of size *s* we must also store O(*s*) weight values, which effectively doubles the space requirement compared to the basic MinHash approach. There is also the added computation of determining the weights. In the baseline approach mentioned above, we count the number of genomes each k-mer in the sketch is present in; this can take O(*n*) time, though this can be minimized by using existing optimized implementations, such as KMC3 (Kokot, Dlugosz, and Deorowicz 2017) or Jellyfish (Marcais and Kingsford 2012).

#### Order MinHash (OMH)

As an alternative to WMH, the consideration of the order of the retained minimal hashes can also filter out spurious matches and prioritize more likely sources for a query sequence. First presented as a method to improve estimation of the edit distance (Marçais, DeBlasio, et al. 2019), an Order MinHash sketch stores the selected *n* hashes in ordered sublists of *L* hashes, in the same order as they occur in the genome, with *n/L* lists making up the sketch. When two sketches are compared using Order MinHash, the algorithm checks which hashes are shared, along with if the shared hashes are in the correct order relative to each other. This method of comparing two sketches means that two sequences that contain the same k-mers but are rearranged versions of each other will have low similarity scores, while non-ordered MinHash would report high similarity. This approach is also more robust to sequencing errors than selecting a single long k-mer spanning the same distance.

As with Weighted MinHash, Order MinHash incurs additional computational overhead. During sketch generation, we must store both the k-mer’s hash and its position in the original sequence, in order to construct the ordered sublists. In addition, during comparisons between two sets of hashes, the relative order of any shared hashes must also be considered, meaning simple set comparison is no longer enough.

A practical limitation of approaches that include ordering in similarity comparison is that they may not be completely suitable for circular genomes, where the relative ordering of k-mers is not possible without assuming that all compared sequences have agreed on the same starting point within the circle. However, with an agreed starting point for the sequence, only sub-lists of k-mers that span this starting point will be affected by the circular nature of the original sequence. With short sub-list lengths, as is the case in OMH, we can limit the impact of this to just a handful of elements in the sketch. This is comparable to index-based approaches not considering the k k-mers that span the circle, or alignment approaches not extending alignments at the end of linear sequences to account for the circle.

#### Minimizer

Minimizers were originally proposed as a sequence compression method (Roberts et al.2004), but have become popular in genomics due to their ability to succinctly represent large sequences. In the most widely used form of a windowed-minimizer, the algorithm slides a window of size *x* over the sequence, and the k-mer with the smallest hash in that window is retained as the minimizer. This is repeated across the entire sequence, and the set of unique minimizers is used as the representation of the full sequence. Unlike MinHash, window-minimizers provide some guarantee on the distance between the retained k-mers in our screen, as this distance is bounded above by twice the window size. For our minimizer-based approach, the window size *w* is computed as the size of the genome (*n*) divided by the desired number of k-mers per genome (*s*), multiplied by a fixed multiplier of two. The reasoning behind this multiplier is quite simple: since the distance between minimizers is uniformly distributed between 0 and the window size *w*, we expect two minimizers per *w* bases of sequence. This means that without the multiplier, we expect 2*s* minimizers across the genome. Consequently, to find *s* minimizers for a genome of size *n*, we simply double the window size. With this change, we expect a minimizer every *w* bases, instead of every *w*/2, and thus a total of *s* minimizers across the genome. This keeps the generated sketch and sample sizes relatively even across the approaches. Computation of a minimizer sketch of a genome of size *n* using window size *w* can be done naively in O(*nw*) by choosing the minimum of the *w* hashes in each of the O(*n*) windows, or in O(*n*) by using an integer representation of the k-mers in the sequence.

### 2.3 Clustering sketches and samples

Using the approaches above, we can construct a screen containing reduced representations of each source genome. Input reads are then compared against all elements of this screen, and their source is predicted based on their similarity to those elements. While this greatly reduces the comparisons necessary to classify a read compared to traditional approaches, they still perform a large number of unnecessary comparisons with genomes with low similarity with the query read. To tackle this, we propose a clustering-based approach to limit the number of comparisons with less-relevant genomes.

To do this, the algorithm first computes a hierarchical clustering of the individual sketches of the input genomes. This groups together similar genomes, whose selected k-mers (and derivative reads) are more likely to be similar. The elements of the screen are then generated as before, using the chosen sketching or sampling technique outlined in the previous section. However, instead of then generating the screen as normal, we can use the generated sketches or samples to populate the calculated clustering tree. Each genome’s reduced representation appears in the leaves of the tree, and the reduced representations are combined within internal nodes of the tree, until the root of the tree contains all the elements of the original screen.

To limit the overhead of this approach to be comparable to those presented above, the algorithm randomly downsamples the original elements of the screen as the tree is constructed from the sketches and samples. This downsampling can be done as a constant factor or by a factor proportional to the height of the tree, depending on the desired total size of the sketch tree. A random downsample could distort set comparison metrics such as Jaccard coefficient estimation, but is less of a concern for this analysis since a single k-mer match is sufficient to explore the children of an internal node. However, extreme downsampling can increase the number of misclassified or unclassified reads as they can remove all shared k-mers.

Read classification is then performed by starting at the root, comparing the input read to the stored representations at each of the children of the root, and then descending into the child with which the read is most similar. This process repeats until the algorithm arrives at a leaf, which is the genome predicted to be the source of the read. If, at any point, no child is found with similarity above the required threshold to the query read, the read is left unclassified. This approach quickly prunes genomes with low similarity to the input read, and focuses on the genomes that are likely to be the source of the read, especially genomes that are either from different but similar organisms, or different assemblies of the same genome.

Using a clustering-based approach can also provide more control over the classification process. Instead of simply classifying reads to a single genome, we can classify reads to a cluster of potential source genomes with high similarity by stopping the classification process before it reaches the leaves. This is similar to the LCA classification approach taken in some index-based approaches (Wood and Salzberg 2014; Marić et al. 2020; Kim et al. 2016), and can be useful in some scenarios. We can also utilize this approach to better understand where and why misclassifications occur, by following the classification of a read down the tree. Doing so can allow us to identify exactly where misclassifications occurred, and allow us to determine how far from the correct genome the predicted genome is.

Hierarchical clustering is computationally expensive, requiring O(*n*^3^) time and Ω(*n*^2^) space to cluster *n* elements. This is in addition to the cost of sketching genomes to input into the clustering and the cost of the initial screen generation. However, the reduction in the number of comparisons performed per read can compensate for this cost. Given a screen of size *x* hashes, and the same screen in this clustered form with downsampling factor *f*, a read will require *x* comparisons against the original screen but O(*x*/*f*) comparisons with the clustered screen. Therefore, with reasonable choice of *f* we can reduce the computation per read classified and yield an overall amortized time savings.

## 3. Results

### 3.1 Metagenomics Classification

Our main experimental results are based on the widely used Culturable Genome Reference (CGR) community of high quality microbial genomes sequenced from the human gut (Zou et al. 2019). From this community, we selected all genomes that were available on RefSeq (as of 12/10/20), giving us 1,310 genomes for our reference database. This community contains several clusters of highly similar genomes that make read classification more difficult (**Table 2, Figure 2**). This difficulty is especially true for approaches that work with reduced representations of the original genomes; unless the differences between these similar genomes are specifically captured, there will be no information available to distinguish between them.

**Figure 2:**
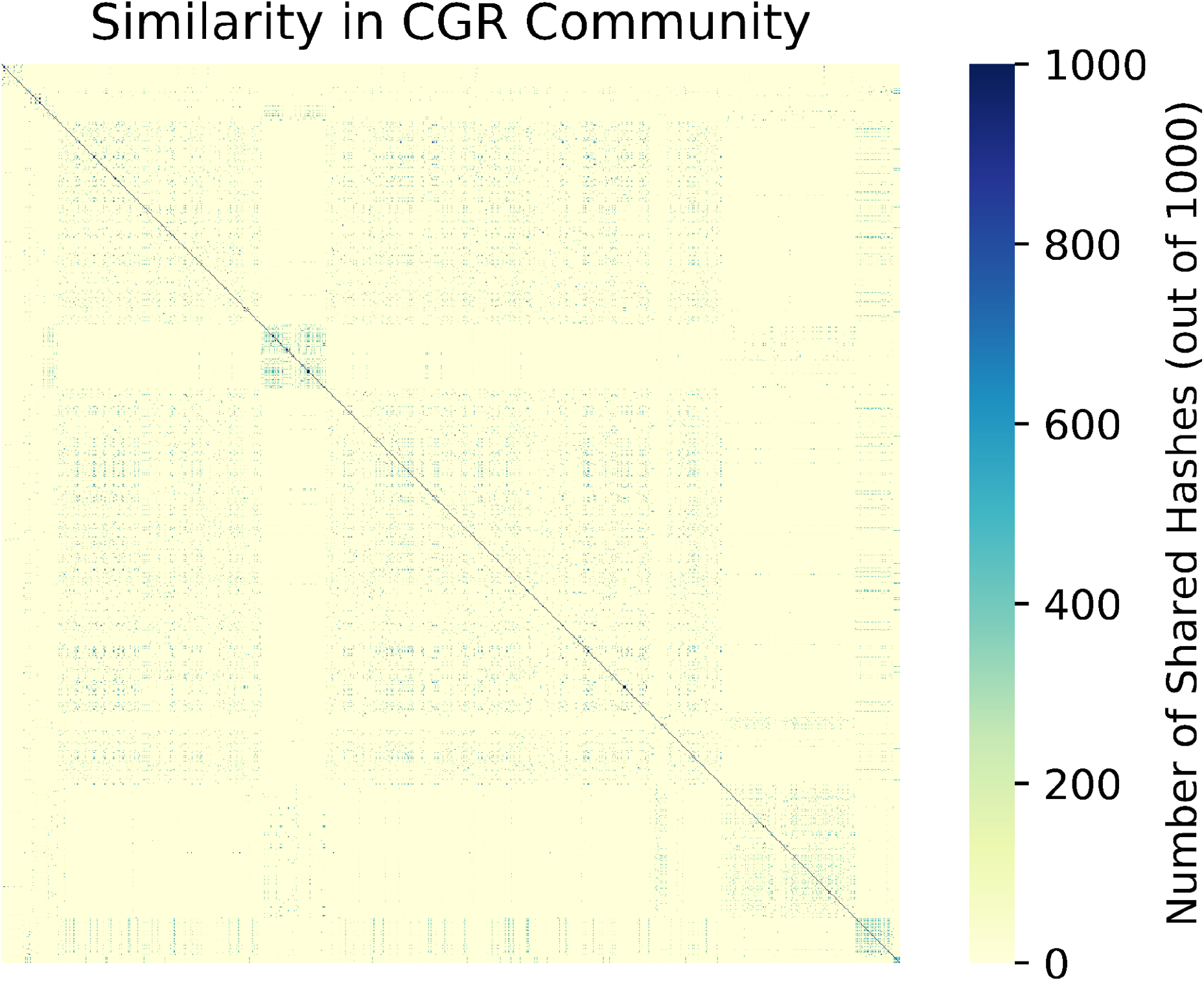
Similarity in the CGR Community. Similarity between the 1310 members of the CGR community, calculated using the number of shared hashes in their Mash sketches. In comparison to the ZYMO + MBARC-26 community (**Supplementary Figure 1**), there are many clusters of high similarity in the CGR community. Genomes within these clusters are very difficult to distinguish between, and contribute to the lower classification accuracy, across all classification approaches, in this community.

**Table 1:**
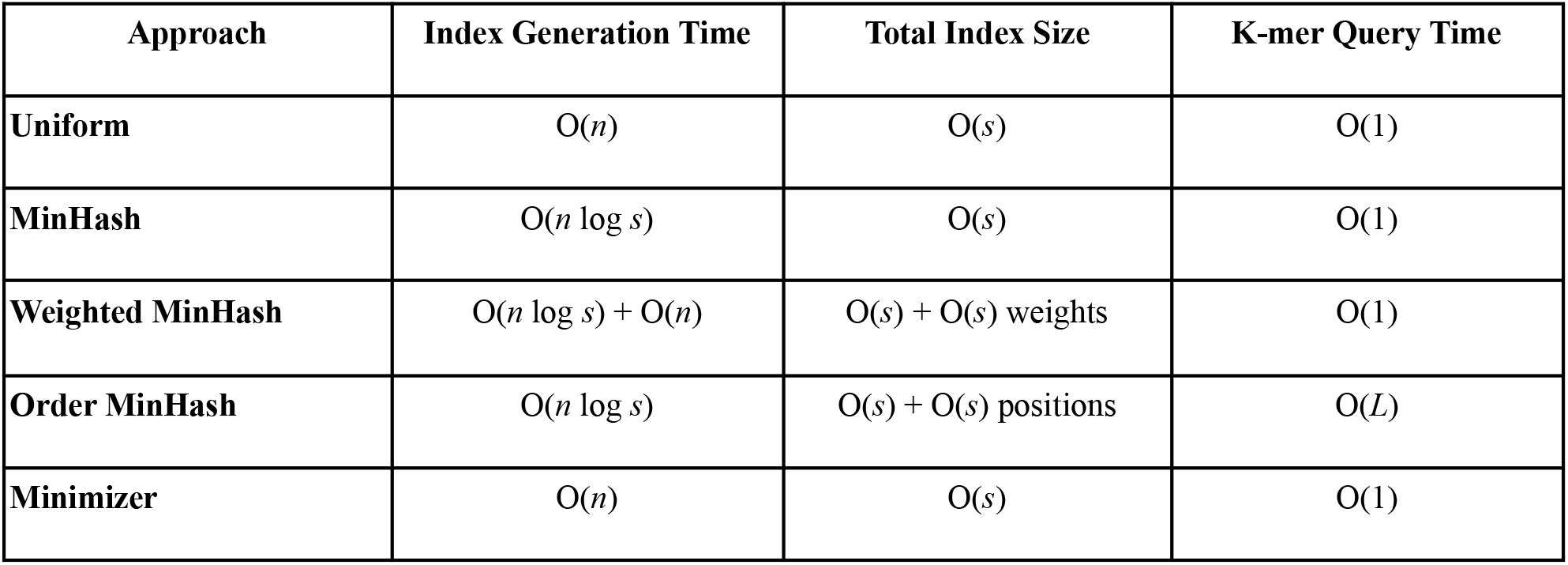
Comparison of sketching and sampling approaches. The theoretical runtimes for generating and querying screens of size *s* generated from a genome of size *n*. The main three approaches (Uniform, MinHash and Minimizer), are largely equivalent in terms of their computational cost. The augmented approaches (Weighted, Order) incur additional overhead, with Order MinHash also involving a more complex query process when comparing two sketches, depending on the choice of size of sublists (*L*). Exact counts of screen sizes and the number of lookups performed during a classification experiment, as well as the overhead of an exhaustive approach, can be found in **Supplementary Table 5**.

| Approach | Index Generation Time | Total Index Size | K-mer Query Time |
| --- | --- | --- | --- |
| **Uniform** | $O(n)$ | $O(s)$ | $O(1)$ |
| **MinHash** | $O(n \log s)$ | $O(s)$ | $O(1)$ |
| **Weighted MinHash** | $O(n \log s) + O(n)$ | $O(s) + O(s)$ weights | $O(1)$ |
| **Order MinHash** | $O(n \log s)$ | $O(s) + O(s)$ positions | $O(L)$ |
| **Minimizer** | $O(n)$ | $O(s)$ | $O(1)$ |

**Table 2:** Overview of the microbial communities.

| Community | Total Sequence | Number of Organisms | Number of genomes with >X% Mash similarity to another genome in the community |  |  |  |  |  |
| --- | --- | --- | --- | --- | --- | --- | --- | --- |
|  |  |  | X = 5% | X = 25% | X = 50% | X = 70% | X = 90% | X = 95% |
| **CGR** | 4.85GBp | 1310 | 1218 | 1189 | 990 | 563 | 315 | 270 |
| **ZYMO +**<br>**MBARC-26** | 170MBp | 34 | 2 | 0 | 0 | 0 | 0 | 0 |

As an example of a simpler community, we also analyze a union of the ZymoBIOMICS Microbial Community Standards (ZYMO) and MBARC-26 (Singer et al. 2016) reference communities, which, when combined, contain 34 microbial and fungal genomes. This community is much easier to classify within, as the genomes are relatively dissimilar (**Table 2, Supplementary Figure 1**), and provides a baseline from which to interpret our results.

For experiments in this work that use simulated reads, we use a simple simulator we developed that allows for simulating reads across a range of read lengths and error models. We chose to use this simulator as we are not trying to exactly represent the error models of a particular technology, and instead are exploring a wider range of read lengths and error rates than currently found in genuine HiFi or ONT reads. A simple simulator allows us to better explore the effects of read length and error rate, and allows us to model performance on data with a range of parameters. More details about this read simulator can be found at https://github.com/arun96/sketching.

For our classification analysis, by default we use simulated PacBio HiFi-like reads that are 10Kb long with 1% error rates, with errors uniformly introduced. We simulate 10x coverage of each genome, yielding 4.8M reads for the CGR dataset, and 165K reads for the ZYMO+MBARC-26 dataset. For the classification, we use screen sizes that target 100 shared hashes with each of these reads, or on average one shared hash every 100bp based on **EQ1**; this generates screens that contain approximately 2% of the k-mers present in the original genomes.

Results across a range of read lengths, error rates and screen sizes, as well as other experimental parameters, are reported later in this section, and presented in **Figure 3**. As no read simulator can perfectly capture the complexity of real data, results on genuine metagenomic sequencing data can be found in section 3.4. Reference implementations, analysis scripts and details on data availability can be found at https://github.com/arun96/sketching.

**Figure 3.**
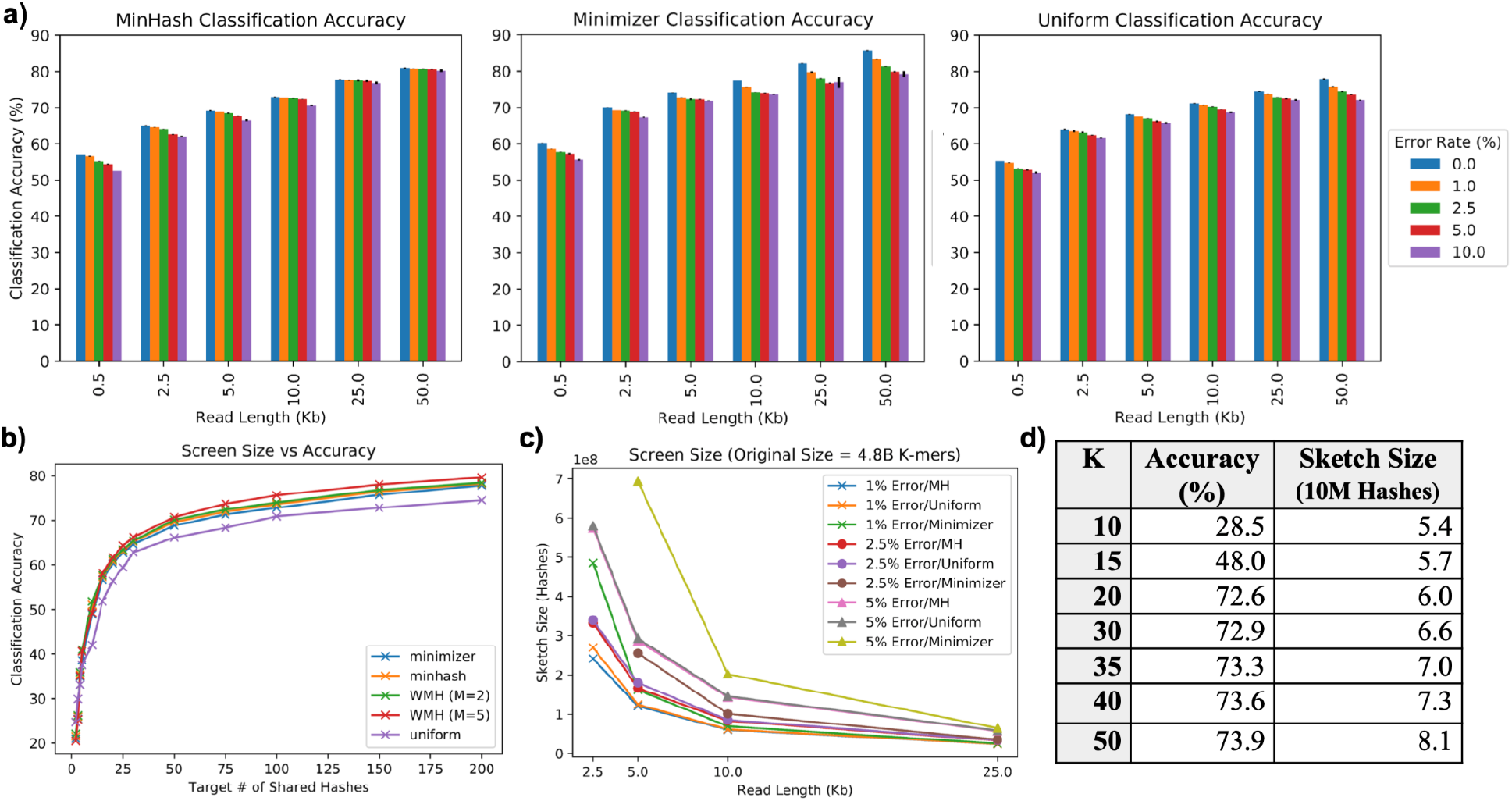
Key results across parameters on the CGR dataset. **(a)** Classification accuracy across a range of read lengths and error rates, averaged over three simulated runs with variance between runs noted on each bar, **(b)** the effect on accuracy of increasing the target number of shared hashes, **(c)** the impact read length and error have on sketch size across our approaches, and **(d)** the accuracy and overhead of a MinHash approach across a range of k values.

#### Classification Experiments

In microbial classification experiments, reads are drawn from a microbial community, and compared against a screen generated from a reference database of known genomes. Under idealized conditions, the database will contain reference genomes from all members of the community, although in practice the community may contain novel species or strains that are not yet characterized leading to poor matches or no matches at all. For simplicity, our simulated reads are drawn from the reference database collection, and reads are then classified against the screen of all genomes. Accuracy is then measured as the fraction of reads correctly classified as being from the true source genome.

For the human gut microbe community, at our default experimental parameters, we see that all our sketching and sampling approaches achieve approximately 71-75% accuracy (**Figure 3**).We observe that around half of the genomes have classification accuracy over 90%, with the overall accuracy lowered by genomes that have high similarity with other members of the community. For example, we find that of the genomes with less than 50% classification accuracy, 90% have another member of the community with which they are at least 70% similar. This implies that for these genomes, we expect 70% of their reads to be very similar to at least one other genome, greatly increasing the chances of each read being misclassified. This is magnified when considering groups of genomes with >95% similarity, a level of similarity high enough to consider the genomes to be of the same species (Jain, Rodriguez-R, et al. 2018). Read classification between such genomes, regardless of which approach is used, often becomes random tie-breaking.

Just how disruptive highly similar genomes are to classification accuracy is visible when classifying reads from the simpler ZYMO+MBARC-26. Here, just two of the 34 members have Mash similarity >1.5% with each other, with those two members having a similarity of just 8% (**Table 2**). In our experiments, these two genomes provide the vast majority of misclassified reads, across read lengths and error rates. Overall, with a simpler community like this, any of these approaches achieve >99% classification accuracy, even with much shorter reads and significantly higher error rates.

The practical consequences of these misclassifications between highly similar genomes depends on the specific downstream analysis used in the experimental scenario. However, it is clear that misclassifications will cause a decrease in precision if reads unrelated to the genomes of interest are mistakenly classified as being of interest, and will cause a decrease in recall if reads of interest are incorrectly classified to genomes filtered out from later analysis.

#### Impact of novel sequences on read classification

In practice, the readset being classified may contain reads from novel species or strains not present in the set of potential source genomes. To model this situation, and to understand where reads from such a novel source genome may go when classified using our approaches, we removed selected genomes from our set of potential sources. We did this for genomes with three different levels of Mash similarity to other members of the community: one genome with no other members with >50% similarity to it, a second with members that it had between 50% and 90% similarity with, and a third with members with greater than 90% similarity to it.

We find that for the first scenario, with no similar genomes in the reference collection, removal from the set of potential source genomes results in the vast majority of its reads remaining unclassified. For the second scenario, which retains some similar genomes but no highly similar genomes, the majority of its reads remain classified, but approximately 25% of its reads are incorrectly classified to one of its most similar counterparts. Finally, for the third scenario with a number of highly similar genomes, the vast majority of its reads are classified to these highly similar genomes, with only a few reads remaining unclassified.

These results are in line with what we expect from a similarity-based classification method, and indicates that the classification of reads from a novel strain not represented in our set of potential source genomes depends on the similarity of the novel strain to the genomes its reads are being compared against.

#### Effect of experimental parameters on read classification

##### Read Length

We see increases in performance as read lengths get longer (**Figure 3a, Supplementary Table 1**), as we have more opportunities for the screen k-mers to match error-free k-mers in the read. Read length also affects the size of the screen, as longer reads mean smaller screens are necessary to achieve the desired number of shared hashes between a read and its source (**Figure 3c**). Conversely, with shorter reads, the screen sizes must be proportionally larger to maintain the similar levels of accuracy.

##### Error Rates

We see decreases in accuracy at high error rates (**Figure 3a, Supplementary Table 1**), as fewer k-mers remain unaffected by error, accompanied by sharp increases in screen size. With an error rate of 1% (as found in PacBio HiFi reads), we estimate that 81% of 21-mers will remain error free, while at an error rate of 5% (as is found in Oxford Nanopore reads) just 34% of the 21-mers will remain error free. This is even more pronounced at error rates close to 10% (as is found in CLR PacBio data and older Oxford Nanopore reads), where just 10% of the 21-mers can be expected to be unaffected by error. As our approach adjusts screen size to compensate for error rate, this results in extremely large screen sizes to compensate for high error rate (**Figure 3c**).

##### Target number of shared hashes

The number of target matches determines how densely the reference genomes are represented, and therefore the size of the screen. Low numbers of target matches result in large numbers of reads being misclassified or unclassified, as there will not be enough detectable similarity with the source genome **(Supplementary Table 2)**. Increasing the number of target matches, and thus the level of similarity between a query read and its source genome, causes sharp improvements in accuracy **(Supplementary Table 2)**. However, there is also a plateau in performance as we increase the target number of shared hashes, as some sets of genomes differ only in a small number of k-mers, and a sketching or sampling approach must draw from exactly those places in order to distinguish between them. We see steady increases in performance when increasing the target number of shared hashes up to 3 shared hashes every 200bp, but performance gains slow beyond this (**Figure 3b**).

##### K

For k set to at least 20 (**Figure 3d**), we find increasing k can result in minimal increases in performance, yet a larger increase in screen size. This is because longer k-mers have a higher chance of being affected by errors, so larger samples/sketches are necessary to ensure a robust number of error free k-mers remain. This was highly pronounced in our results, e.g. the step up from k=30 to k=50 came with only a 1.3% increase in performance but a 22% larger screen. As we saw similar performance between 20 ≤ k ≤ 50, we used k = 21 across our other experiments, as it provided the specificity necessary while keeping screen sizes small.

##### Weight

The addition of weight to traditional MinHash results in a slight increase in performance across read lengths and error rates. This is expected, as the discriminative k-mers now contribute more to the score and help break ties. Including a multiplier has a similar effect and more heavily weighting unique k-mers results in correctly breaking even more ties, resulting in another slight increase in performance. The addition of weight saw a 0.5% increase in classification accuracy over unweighted MinHash (72.7% vs 72.2%), with the inclusion of a multiplier of 5, 10 or 15 seeing a further 0.1% increase in performance (**Supplementary Table 4**).

##### Order

Including order in a MinHash approach has a minimal impact on classification accuracy (**Supplementary Table 4**), giving a 0.2% increase compared to MinHash (72.4% vs 72.2%). Order MinHash was initially proposed as a metric for estimating edit distance, and is most beneficial when determining the similarity between rearranged strings that cannot be distinguished by an unordered MinHash. However, with read classification from this large set of microbial genomes, such rearrangements are not common.

##### Cluster Downsampling Rate

Using MinHash screens, we compared the accuracy across three approaches to clustering: (1) screens downsampled by a constant factor; (2) screens downsampled based on their height in the sketch tree; and (3) screens that are not downsampled at all. We find that constant factor downsampling approaches, with factors *f* = 2 and *f* = 4, maintain a good degree of accuracy (71.1% and 70.1% respectively, compared to the 72.8% accuracy with MinHash), while keeping the number of comparisons similar to or less than the original MinHash approach. Height-based downsampling approach results in a sharp drop in accuracy (62.1%), as the screens near the root of the tree are downsampled to the point where discriminative k-mers are lost.

#### Analysis of clustering-based approach

Examining the output of the clustering-based approaches with zero or constant factor downsampling, we find that the majority of misclassified reads are first misclassified only a few levels from the leaves of the tree. This is in line with the misclassification rates between highly similar genomes; we find that over 90% of the misclassified reads are misclassified to a genome that is at least 90% similar to the true source genome, and about 80% are misclassified to genomes that are at least 95% similar. These misclassifications occur lower down the clustering tree, where these similar genomes are close to each other and classification is forced to choose one path or the other.

This can be seen by looking at the number of reads that are only misclassified close to the end of the classification process. With a constant downsampling factor of *f* = 2, we find that 80% of misclassified reads are only misclassified at the very last level of the tree. These reads come from the many small groups of genomes with high similarity. We can expand this analysis further to find that more than 90% of misclassified reads are only misclassified within the last three steps of their path to a leaf; these reads come from slightly larger groups of highly similar genomes. The remaining misclassified reads mostly come from the few larger clusters of highly similar genomes, with a few stemming from similarity caused by randomly downsampling the generated sketches or samples. When the downsampling rate is increased to *f* = 4, we see slight increases in the number of reads misclassified further up the tree, but the overall pattern holds.

We can also explore these clustering-based results to highlight the small margins that cause misclassifications.With a constant downsampling factor *f* = 4, just over 75% of all misclassified reads result from incorrectly breaking a tie, and a further 15% from an incorrect source having just one more shared hash with the read than the true source. Decreasing this downsampling factor to *f* = 2 drops the latter to 8%, with incorrect tie breaks accounting for 87% of all incorrect classification decisions. This means that with downsampling factors *f* = 2 and *f* = 4, 90% and 95% of misclassifications respectively come down to tie breaks or a single extra shared hash across the entire read.

An alternative approach to help avoid misclassifications based on incorrectly resolving ties is to end classification when a tie is encountered, and report that the source genomes are found in the subtree below the internal node where the tie occured. As ties are the primary cause of incorrectly classified reads, and mostly occur near the leaves (thus meaning the subtree rooted at them is small), such an approach would greatly narrow down the source of a large number of previously misclassified reads without incorrectly associating a read to a single wrong genome. Applying this method to our previous results, we find the leaf representing the predicted source of the read is in the subtree below where classification is stopped 90% of the time, even with a downsampling factor of *f* = 4. This method is commonly utilized in index-based read classification approaches, and the choice to use a clustered screen opens the door to this form of classification, which is suitable for applications where narrowing the source of a read to a small group of highly similar potential source genomes instead of a single genome is sufficient.

It is worth mentioning that the pattern of the majority of misclassifications occurring lower down in the tree does not hold when using height-based downsampling. In addition to the drop in accuracy discussed in the previous section, we observe high numbers of misclassification in the initial levels of the clustering tree, as the elements there are downsampled to the point where distinction between even slightly similar elements is difficult.

### 3.2 Host Contaminant Detection

The goal of the contaminant detection and classification is, for a given read set, to distinguish between reads that come from organisms or sequences of interest, and reads that are from potential contaminants. For our experiments, we considered human reads simulated from GRCh38 (Schneider et al. 2017) and mixed with contaminant reads drawn from a selected microbial community, and classified against a screen containing both the human and microbial genomes. The HiFi-like sequence reads were simulated as above using 10Kb reads at 10x coverage with 1% error. Accuracy is measured as the fraction of human and microbial reads correctly identified as being of interest or as being a contaminant, while also measuring the fraction of contaminant reads that are correctly classified to their source genome.

When using the CGR community as the source of contaminant reads, we find all the sketching and sampling approaches to be successful at distinguishing between microbial and human reads. Across all approaches, >99% of all human reads are correctly distinguished from microbial reads and classified to the chromosome they are drawn from. We also observe that very few microbial reads are misclassified as human, with over 99% correctly identified as being contaminants. This is not unexpected; human and microbial genomes are quite dissimilar, and therefore the sketches or samples of the sequences will also be dissimilar, making read classification successful in nearly all cases.

This lack of similarity can be highlighted by comparing 1Mb regions of the human genome to each of the 1,310 microbial genomes in the contaminant community using Mash. This results in approximately 3.8 million pairwise comparisons, and of these, we find that less than 7,000 of these comparisons share a hash between their sketches. Within these pairs, the level of similarity is low, with very few of these pairs sharing more than 0.5% of the hashes that make up their Mash sketches. This low similarity explains the ease with which we can distinguish most human and contaminant reads, but also explains the small number of the reads that remain incorrectly identified, as these reads may be drawn from these small regions of similarity.

Despite the dissimilarity between the human and contaminant genomes, it is worth highlighting that we are able to distinguish between these sequences with high accuracy while storing just 2% of the original k-mers. We see similar accuracy while storing as little as 1% of the original k-mers, with a slight decrease to 98% when storing 0.2% of the k-mers and 96% when storing 0.1% (**Supplementary Table 3**). Below this threshold, accuracy starts to drop sharply; when storing just one of every 2000 k-mers, we are able to differentiate between 89% of human and microbial reads, and 60-70% when we further halve the number of stored k-mers (**Supplementary Table 3**).

After distinguishing between human reads and contaminants, we then attempt to classify the contaminant reads to the exact source genome. The results match what we present in the classification experiments; approximately 75% of the contaminant reads are mapped to the correct genome, with the misclassified reads coming from the genomes discussed in the previous section. We also classify human reads to the chromosome they are drawn from, and are able to do this with >95% accuracy.

### 3.3 Comparison to existing tools

To evaluate the performance of the sketching and sampling approaches, we also tested several widely used approaches for read classification on the same dataset and experimental settings. Versions of all tools used can be found in **Supplementary Note 1**.

In order to compare these existing tools to both our reference implementations of the sketching and sampling approaches and to each other, these comparisons are done using accuracy instead of runtime. This is done to simplify the comparison between tools with different levels of optimization, and allows us to focus simply on the ability of these approaches to correctly perform the task at hand. However, some details on the overhead of the index-based approaches can be found in **Supplementary Table 6**, and comparisons between the runtime of selected index- and alignment-based approaches can be found in **Supplementary Table 7**. These results highlight the wide range in practical requirements between even highly similar approaches, further emphasizing why we have chosen to focus on accuracy as the key metric for comparison.

As before, accuracy in classification experiments is measured as the number of reads correctly classified as from the microbial genomes they were drawn from, and accuracy in contaminant detection experiments is measured as the number of human and microbial reads identified as human or from any microbial genome respectively.

#### Alignment-based

To test the effectiveness of alignment-based approaches to read classification, we test **Minimap2** (H. Li 2018) and **Winnowmap** (Jain, Rhie, Zhang, et al. 2020). Minimap2 uses query minimizers as seeds for the alignment, while Winnowmap2 adds a preprocessing step to downweight repetitive minimizers to reduce the chance of them being selected. In both approaches, we align our read sets against the genomes of the selected community, and calculate the predicted source of the read as the sequence to which it is mapped. For microbial classification, we find that both these tools perform slightly better than our MinHash and minimizer based approaches. Compared to an accuracy of 77% and 79% in our MinHash and minimizer approaches with two shared hashes every 100bp, Minimap2 and Winnowmap both achieve an accuracy of 81% (**Table 3**). Both alignment approaches achieve low accuracy on the same genomes that our sketching and sampling approaches struggle on; namely, genomes with high-similarity relatives in the community. These misclassifications are amplified at higher error rates, where the ability to distinguish between similar genomes is reduced. For contaminant detection, both tools are able to correctly distinguish 99.5% of the human and contaminant reads at a 1% sequencing error rate, and over 98% at higher error rates.

**Table 3:**
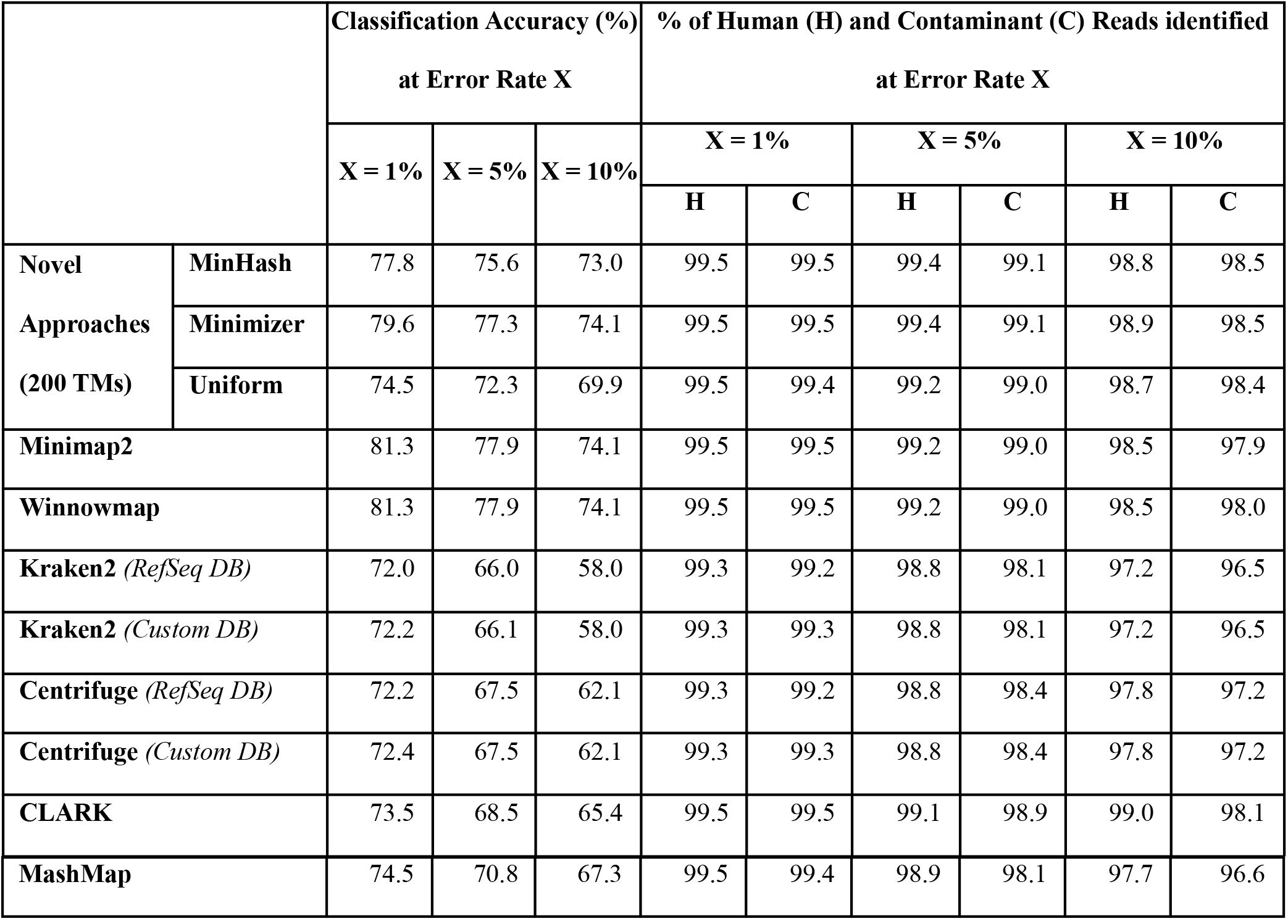
Performance of existing tools. For genome-level classification accuracy, we find that alignment based methods perform best, due to their ability to compare against the entire sequence instead of a reduced or indexed form, allowing them to identify minute differences between highly similar genomes. Index-based approaches struggle to perform genome-level classification between highly similar genomes, with a significant number of reads being classified only to a lowest common ancestor of several possible source genomes. All tools perform similarly in contaminant detection, with this task less affected by higher error rates.

#### Index-based

**Kraken2** (Wood, Lu, and Langmead 2019) and **Centrifuge** (Kim et al. 2016) use a preprocessed index of shared k-mers or compressed genomes respectively to determine the source of a query sequence. Each k-mer in the query sequence is classified to an element in the index, and we determine the source of the query as the element to which a plurality of k-mers are assigned.When using both a pre-built RefSeq database and a custom database built over our test community, we find that genome level identification is difficult between the highly similar members (**Table 3**). We observe large numbers of misclassifications between reads from these similar genomes, as well as classifying many of these reads only to higher taxa, and not to one of the specific genomes. For genomes without similar members in the community, the majority of their reads are correctly classified, giving Kraken2 and Centrifuge an overall classification accuracy of 72% with 1% error reads. At higher error rates, this performance drops sharply, with more reads left unclassified due to a lack of matched k-mers to the generated index. When distinguishing between human and microbial reads, both methods are able to correctly identify >95% of the reads, even at high error rates.

#### CLARK

(Ounit et al. 2015) uses a pre-compiled list of discriminative k-mers for the community it is indexing, and performs classification based on query similarity to this list. While there are still misclassifications and unclassified reads at rates comparable to other tools, CLARK’s use of discriminative k-mers slightly reduces the impact of highly similar genomes in the community, allowing it to identify the few differences between them, achieving a classification accuracy of 73.5% with 1% error reads, and making it more resilient against misclassification at higher error rates (**Table 3**). For contaminant detection, CLARK is also able to distinguish over 97% of both human and microbial reads across a range of error rates.

#### Sketching-based

**MashMap** (Jain, Dilthey, et al. 2018) computes alignments by estimating k-mer based Jaccard similarity between query sequences with MinHash sketches. We find that MashMap performs worse than classic alignment-based approaches, and similarly to our MinHash approaches, with 74.5% classification on 1% error reads and steady decreases at higher error rates. Alignment boundaries in MashMap are determined through the Jaccard similarities of sketches. As a result, just as in the MinHash approach, it is susceptible to misclassifications between highly similar genomes. For contaminant detection, MashMap is able to distinguish more than 96% of the human and microbial reads, even at higher error rates (**Table 3**).

### 3.4 Analysis of genuine metagenomics sequencing data

To test the accuracy of our approaches on real sequencing data, we analyzed PacBio HiFi reads from the Human Gut Microbiome Pooled Standards (“Data Release: Human Microbiome Samples Demonstrate Advances in HiFi-Enabled Metagenomic Sequencing” 2021) with the CGR community database. For this analysis, we used one omnivore and one vegan dataset, with 1.79M and 1.90M reads respectively of length ~10Kb and median quality of ~Q40. We first align these reads to the CGR community database using Minimap2, and find that 78% of the reads from the vegan dataset and 85% of the reads from the omnivore dataset align at all. Of these alignments, 56% of reads from the vegan dataset and 63% of reads from the omnivore dataset have alignments that span >50% of the read (**Table 4**), with this fraction dropping to 44% and 48% respectively when looking for alignments that span >90% of the read length (**Supplementary Figure 2**). For reads with alignments to multiple genomes, we take the sequence with the longest alignment to be their predicted source. This is less of a concern for reads with longer alignments, which we find are less likely to be mapped to multiple genomes.

**Table 4:** Performance on real sequencing data. Comparison of the classification of our sketching and sampling approaches against Kraken2 and Minimap2 classifications across two PacBio HiFi Gut Microbiome datasets. The addition of a threshold requiring that >50% of a read is aligned seems to remove a number of more spurious or insignificant calls, increasing concordance between Minimap2 and the other benchmarked approaches.

|  |  | Total number of reads classified | Number of reads classified to multiple genomes | Number of unclassified reads | Number of reads with same prediction as Minimap2 |  |
| --- | --- | --- | --- | --- | --- | --- |
|  |  |  |  |  | No Threshold | >50% of read aligned |
| Vegan<br>(1.90M Reads) | Minimap2 (No threshold) | 1,490,713<br>(78.3%) | 485,233<br>(25.4%) | 413,446<br>(21.7%) | N/A |  |
|  | Minimap2 (>50% of read aligned) | 1,069,306<br>(56.1%) | 23,199<br>(1.2%) | 834,853<br>(43.9%) | N/A |  |
|  | Kraken2 | 1,399,341<br>(73.5%) | 562,744<br>(29.6%) | 504,818<br>(26.5%) | 827,572 | 816,331 |
|  | MinHash | 1,032,396<br>(54.2%) | 261,732<br>(13.7%) | 871,763<br>(45.8%) | 890,766 | 821,556 |
|  | Minimizer | 1,029,996<br>(54.1%) | 262,514<br>(13.8%) | 874,163<br>(45.9%) | 891,388 | 820,493 |
|  | Uniform | 1,021,555<br>(53.6%) | 264,867<br>(13.9%) | 882,604<br>(46.7%) | 883,465 | 819,017 |
| Omnivore<br>(1.79M Reads) | Minimap2 (No threshold) | 1,530,795<br>(85.4%) | 490,501<br>(27.4%) | 261,351<br>(14.6%) | N/A |  |
|  | Minimap2 (>50% of read aligned) | 1,144,452<br>(63.8%) | 24,271<br>(1.3%) | 647,694<br>(36.2%) | N/A |  |
|  | Kraken2 | 1,442,671 | 578,113 | 349,475 | 855,670 | 835,888 |
|  |  | (80.5%) | (32.2%) | (19.5%) |  |  |
|  | **MinHash** | 1,111,102<br>(62.0%) | 275,344<br>(15.4%) | 681,044<br>(38.0%) | 915,634 | 873,901 |
|  | **Minimizer** | 1,105,356<br>(61.7%) | 278,654<br>(15.5%) | 686,790<br>(38.3%) | 916,998 | 872,955 |
|  | **Uniform** | 1,098,244<br>(61.3%) | 281,745<br>(15.7%) | 693,902<br>(38.7%) | 911,554 | 871,675 |

We then classify these reads using our sketching and sampling approaches against a generated screen of the CGR community, built for 10Kb, 1% error reads and 100 shared matches per read. We consider a read to be classified if it has at least 5 shared hashes with a genome, and unclassified if it does not share 5 hashes with any genome. With this threshold, approximately 60% of all reads are classified in each of the approaches, with approximately 25% of the classified reads tied between multiple sources (**Table 4**).

We compare these classification results against the alignments generated with Minimap2. Our classification results agree with approximately 60% of the reads classified by Minimap2 with no minimum alignment length, and approximately 76% of reads who have an alignment >50% of their read length. (**Table 4**). This increase in consistency is expected, as this threshold limits Minimap2 classification to reads that share a significant amount of sequence with potential source genomes, and are therefore more likely to share a significant amount of similarity with elements in the screen. Without this threshold, some reads are classified based on small, potentially unreliable, regions of alignment, and any similarity with elements in the screen must come from shared hashes drawn from these short aligned regions; such reads are likely to be misclassified between multiple low scoring genomes, or not classified at all. An alternate threshold would be to require the alignment to be over a particular length (e.g. 5Kb), but the variance in read length causes this to be skewed against well-aligned shorter reads (**Supplementary Figure 3**). Investigating the classifications that still do not agree with Minimap2, we see that almost 90% of these reads are instead classified to genomes that are >95% similar to Minimap2’s predicted source.

We also classified these reads using Kraken2, with the predicted source of a read being the sequence to which a plurality of its k-mers are assigned. We find that approximately 77% of reads are classified by Kraken2, but approximately 40% of classified reads are only classified to a lowest common ancestor (LCA) instead of a single genome; we count these reads as classified to multiple genomes. We find that Kraken2’s classification results agree with approximately 56% of the reads classified by Minimap2 with no alignment threshold, and approximately 75% of the reads classified by Minimap2 with the >50% read length alignment threshold (**Table 4**). As expected, Kraken2 classified very few reads that fall below the 50% read length alignment threshold Minimap2 alignments, leaving those reads unclassified or only classified to a LCA; the majority of Kraken2’s classifications being reads that have significant similarity to a single genome. The remaining reads that do not agree with Minimap2 are either classified only to a LCA, or classified to genomes that are highly similar to Minimap2’s predicted source. As with the sketching and sampling approaches, we find that almost 90% of these reads are classified to genomes that are >95% similar genome-wide. The remaining mis-classified reads originate from localized regions of the target genome showing high similarity to other genomes.

## 4. Conclusions and Discussion

In this work, we presented and analyzed a range of sketching and sampling approaches for read classification, designed to reduce the space and time overhead for accurate classification across large collections of genomes. Overall, we find sampling and sketching are highly effective compared to index-based approaches, and are within a few percent accuracy of alignment-based approaches. Alignment-based approaches have the advantage that they can assess the entire input sequence, although this increases runtime. Minimizers generally lead to improved accuracy over MinHash-based approaches, chiefly because there is a stronger guarantee on the distance between selected k-mers. Among MinHash-based techniques, weighted MinHash enabled modest but measurable improvements while Ordered MinHash enabled minimal performance gains. All approaches correctly distinguished reads from dissimilar genomes but struggled with the classification of reads from highly similar genomes.

Sketching and sampling approaches are able to perform well, however there are still scenarios where these approaches are challenged. The current methods are best suited for longer, low-error reads, and incur a higher footprint and decreased performance when classifying shorter, higher error rate reads. Consequently, a major need for future work is the continued development of sketching and sampling techniques better suited for high error rate environments. This includes the use of approaches such as gap k-mers (Ghandi, Mohammad-Noori, and Beer 2014) to increase error tolerance, or the use of more auxiliary information, such as pre-computed indexes of unique k-mers (Zhu et al. 2020) or augmented MinHash or minimizer-based methods (Ekim, Berger, and Chikhi 2021; Durbin, n.d.), to distinguish between similar sequences.

## Abbreviations

MH: MinHash
WMH: Weighted MinHash
OMH: Order MinHash
CGR: Culturable Genome Reference
LCA: Lowest Common Ancestor

## Declarations

### Ethics approval and consent to participate

Not applicable.

### Consent for publication

Not applicable.

### Availability of data and materials

All code used for this project, including reference implementations and all analysis or benchmarking scripts, can be found at https://github.com/arun96/sketching. All data used in this work has been cited, and links to each dataset can be found at https://github.com/arun96/sketching#data-and-code-availability.

### Authors’ contributions

Software implementation and computational experiments and analysis were performed by AD, under the supervision of MCS. AD and MCS wrote this manuscript. All authors read and approved the final manuscript.

## Acknowledgements

We would like to thank Benjamin Langmead, Daniel Baker and Bohan Ni for their help and discussions.

## Authors’ information

AD is a PhD student in the Department of Computer Science at Johns Hopkins University, Baltimore, MD, USA.

MCS is a Bloomberg Distinguished Professor of Computer Science and Biology at Johns Hopkins University, Baltimore MD, USA.

## Supplementary Materials

**Supplementary Note 1.**
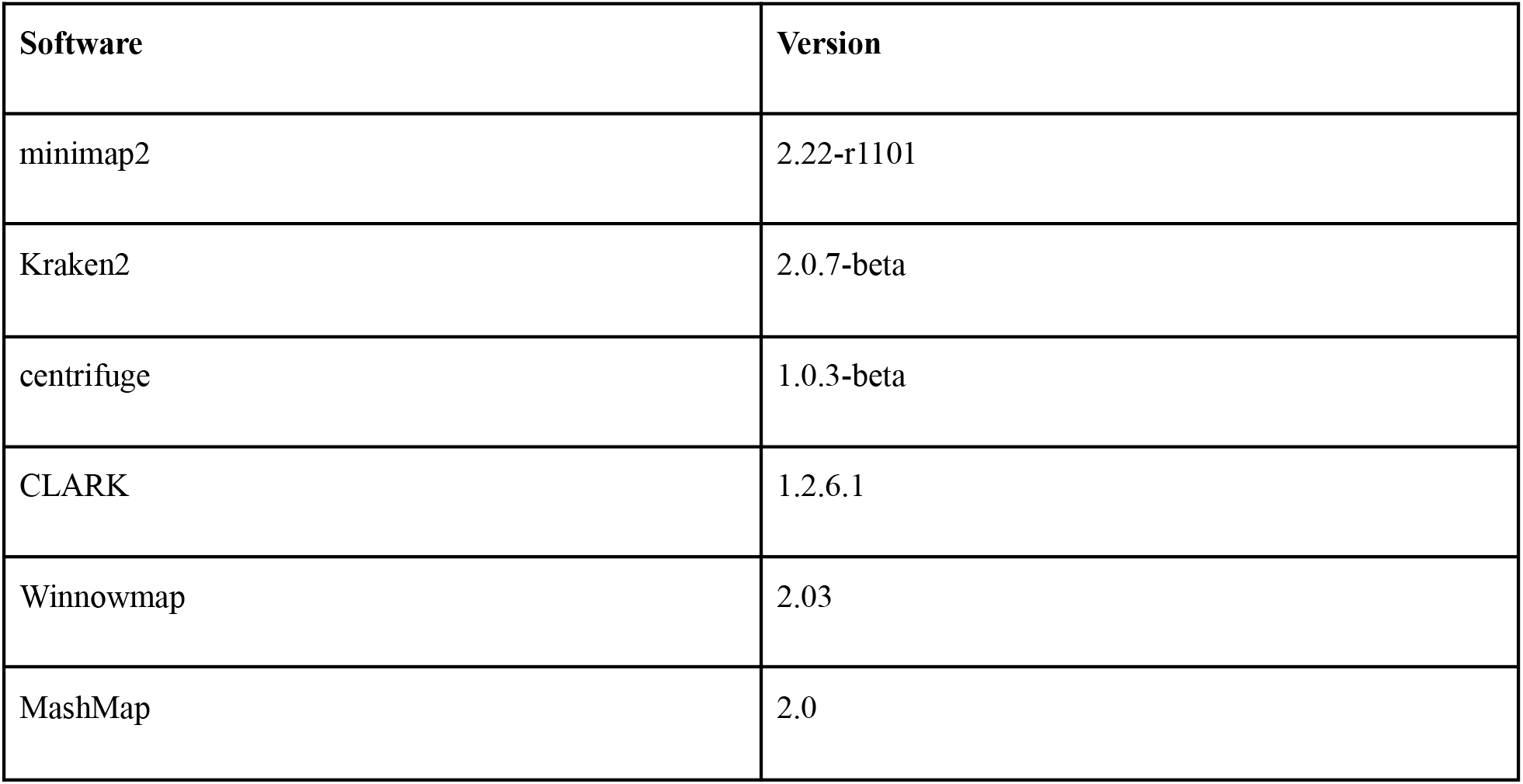
Versions of tools used.

**Supplementary Figure 1:**
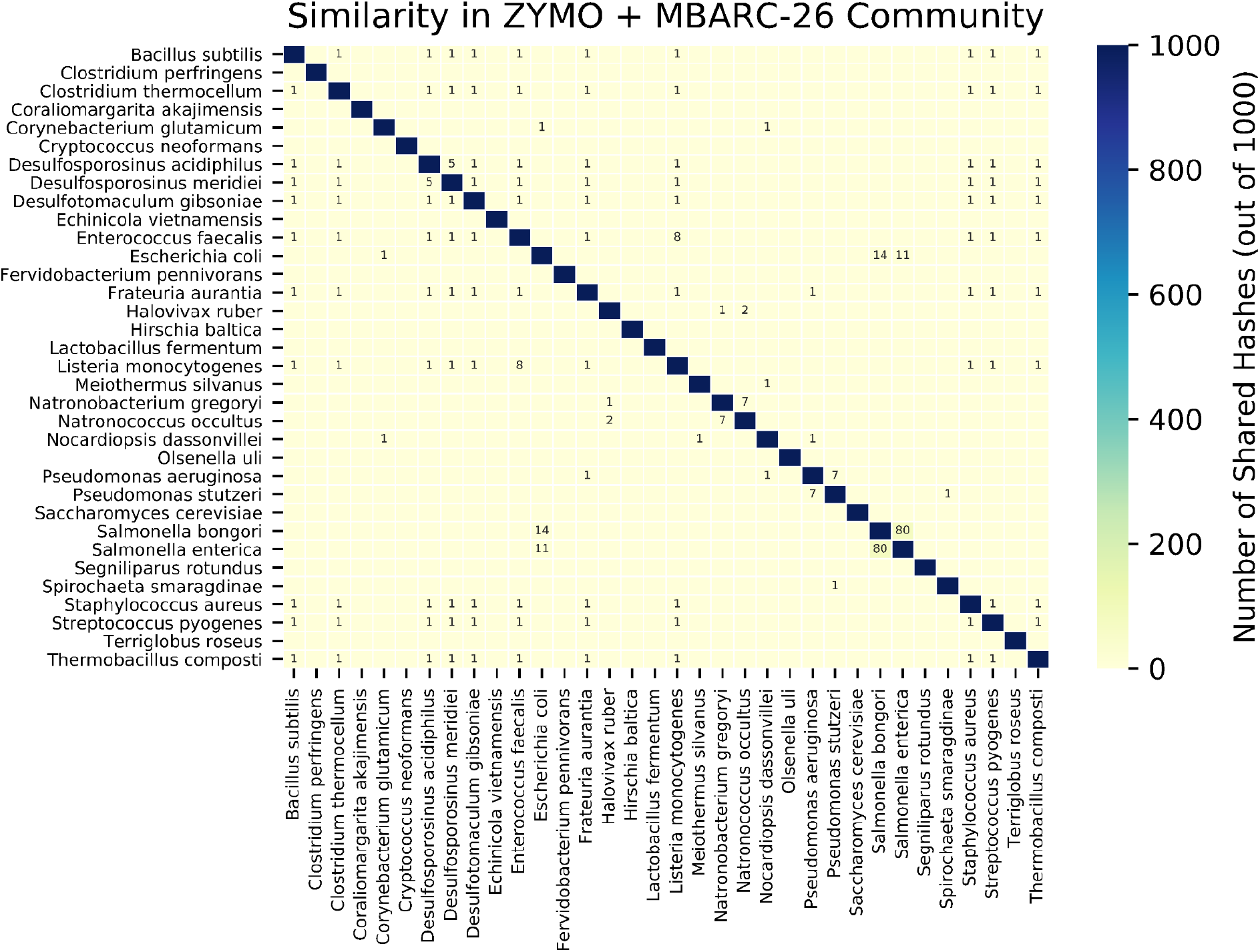
Similarity in the ZYMO + MBARC-26 Community: Using Mash, we measured the similarity between each of the 34 members of the community. Each genome is sketched down to 1000 hashes, and these 1000 hash sketches are compared. All pairs of genomes that share at least one hash are shown on the plot above, with all genomes sharing 1000 hashes with themselves (as seen on the diagonal). The two most similar genomes are two strains of Salmonella, with 8% similarity, but only two other pairs have similarity over 1%. The low level of similarity between members of this community means classification is a much easier task.

**Supplementary Figure 2:**
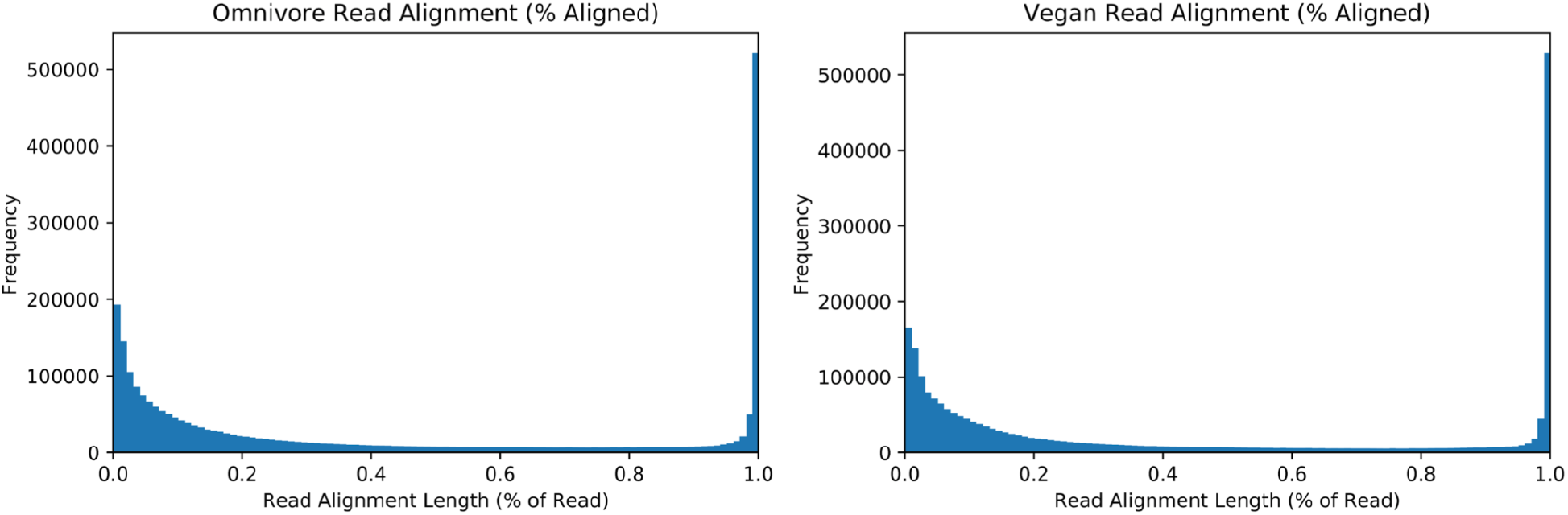
Minimap2 alignment lengths, as a percentage of read length, in two genuine metagenomics sequencing datasets: Distribution of Minimap2 read alignments for genuine PacBio HiFi reads from an omnivore (left) and a vegan (right) to the CGR dataset. We observe a bimodal distribution, with the vast majority of reads either having only small sections aligned (<20%), or being almost completely aligned (>95%), and only a few reads falling in the middle.

**Supplementary Figure 3:**
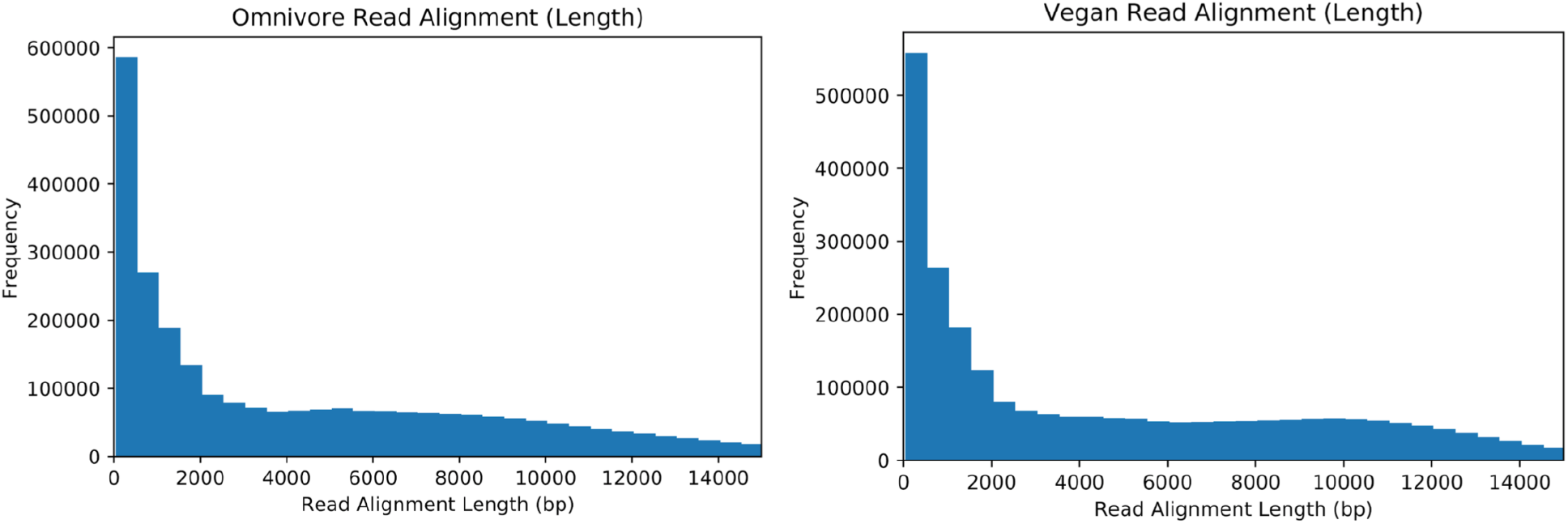
Minimap2 alignment lengths in two genuine metagenomics sequencing datasets: Distribution of Minimap2 read alignments for genuine PacBio HiFi reads from an omnivore (left) and a vegan (right) to the CGR dataset. Unlike **Supplementary Figure 2**, using alignment lengths instead of the fraction of the read aligned allows varying read lengths to skew our analysis. Longer reads with a small fraction aligned may have longer alignments than fully aligned, shorter reads, despite the latter being classified with more certainty.

**Supplementary Table 1:** Microbial Classification Accuracy. Classification accuracy across a range of read lengths and error rates, when using the three main sketching and sampling approaches (MinHash, Minimizer and Uniform).

|  | Read Lengths (Kb) |  |  |  |  |  |  |  |
| --- | --- | --- | --- | --- | --- | --- | --- | --- |
| Read Error<br>(%) |  | 0.5 | 2.5 | 5.0 | 10.0 | 25.0 | 50.0 | MinHash |
|  | 0.0 | 57.1 | 65.0 | 69.0 | 72.8 | 77.5 | 80.8 |  |
|  | 1.0 | 56.6 | 64.6 | 68.9 | 72.8 | 77.6 | 80.8 |  |
|  | 2.5 | 55.1 | 64.1 | 68.6 | 72.7 | 77.8 | 80.8 |  |
|  | 5.0 | 54.5 | 62.7 | 67.8 | 72.4 | 77.7 | 80.7 |  |
|  | 10.0 | 52.5 | 62.0 | 66.9 | 70.8 | 77.2 | 80.6 |  |
| Read Error<br>(%) | 0.0 | 60.1 | 70.0 | 74.0 | 77.4 | 82.1 | 85.8 | Minimizer |
|  | 1.0 | 58.7 | 69.2 | 72.7 | 75.6 | 79.8 | 83.3 |  |
|  | 2.5 | 57.8 | 69.0 | 72.1 | 74.2 | 78.0 | 81.2 |  |
|  | 5.0 | 57.1 | 68.9 | 72.3 | 74.0 | 76.7 | 79.8 |  |
|  | 10.0 | 55.6 | 67.4 | 71.9 | 73.6 | 78.6 | 80.1 |  |
| Read Error<br>(%) | 0.0 | 55.3 | 63.9 | 68.1 | 71.1 | 74.7 | 77.9 | Uniform |
|  | 1.0 | 54.8 | 63.5 | 67.6 | 70.9 | 73.7 | 75.9 |  |
|  | 2.5 | 53.3 | 63.1 | 67.1 | 70.5 | 72.9 | 74.8 |  |
|  | 5.0 | 52.8 | 62.4 | 66.4 | 69.5 | 72.6 | 73.7 |  |
|  | 10.0 | 52.1 | 61.9 | 66.0 | 68.9 | 72.1 | 72.2 |  |

**Supplementary Table 2:** Effect of sketch size on microbial classification accuracy. Classification accuracy on the CGR dataset using sketching and sampling approaches, across a range of target matches. We observe steady decreases in genome-level classification accuracy as the number of target matches decreases.

| # Target Matches | Microbial Classification Accuracy |  |  |  |  |
| --- | --- | --- | --- | --- | --- |
|  | MinHash | WMH (W = 2) | WMH (W = 5) | Minimizer | Uniform |
| **1** | 17.1 | 17.2 | 17.5 | 16.9 | 18.9 |
| **2** | 21.0 | 21.5 | 22.1 | 20.5 | 24.7 |
| **3** | 25.4 | 25.9 | 26.2 | 25.3 | 29.9 |
| **4** | 34.9 | 35.4 | 35.9 | 35.1 | 33.1 |
| **5** | 38.6 | 40.6 | 41.0 | 40.6 | 37.5 |
| **10** | 48.9 | 51.0 | 51.8 | 49.2 | 42.0 |
| **15** | 56.7 | 57.3 | 57.6 | 58.1 | 51.8 |
| **20** | 60.3 | 60.8 | 61.2 | 61.7 | 56.4 |
| **25** | 62.7 | 63.1 | 63.4 | 64.3 | 59.4 |
| **30** | 64.6 | 65.2 | 65.4 | 66.2 | 62.8 |
| **50** | 68.8 | 69.6 | 70.1 | 70.7 | 66.1 |
| **75** | 71.3 | 71.9 | 72.4 | 73.7 | 68.3 |
| **100** | 72.8 | 73.6 | 74.0 | 75.6 | 70.9 |
| **150** | 75.7 | 76.4 | 76.8 | 78.0 | 72.8 |
| **200** | 77.8 | 78.2 | 78.4 | 79.6 | 74.5 |

**Supplementary Table 3:** Effect of sketch size on contaminant detection. Percentage of human (H) or “contaminant” microbial (C) reads correctly identified as being of interest or from contaminants respectively, across a range of sketching and sampling approaches. We find that accuracy remains relatively constant until the number of target matches, and therefore the sketch size, drops below 0.2-0.3% of the original k-mers.

|  | % Human (H) and Contaminant (C) Reads Identified |  |  |  |  |  |  |  |  |  |
| --- | --- | --- | --- | --- | --- | --- | --- | --- | --- | --- |
|  | MinHash |  | WMH (W = 2) |  | WMH (W = 5) |  | Minimizer |  | Uniform |  |
| # Target Matches | H | C | H | C | H | C | H | C | H | C |
| 1 | 45.5 | 42.1 | 46.0 | 42.2 | 46.3 | 42.5 | 46.1 | 43.1 | 45.6 | 42.7 |
| 2 | 59.4 | 56.3 | 60.4 | 56.7 | 60.8 | 57.0 | 63.2 | 60.8 | 61.0 | 60.5 |
| 3 | 72.6 | 70.9 | 73.1 | 71.4 | 73.4 | 71.6 | 75.0 | 73.7 | 72.4 | 70.4 |
| 4 | 83.2 | 82.4 | 83.8 | 82.6 | 84.1 | 82.8 | 84.5 | 83.1 | 82.6 | 80.3 |
| 5 | 89.4 | 88.5 | 90.5 | 88.9 | 90.8 | 89.0 | 90.8 | 89.0 | 88.6 | 88.1 |
| 10 | 96.1 | 95.2 | 96.4 | 95.3 | 96.4 | 95.3 | 96.6 | 95.3 | 95.4 | 94.9 |
| 15 | 97.5 | 96.7 | 97.6 | 96.8 | 97.7 | 96.9 | 97.8 | 96.9 | 97.0 | 96.5 |
| 20 | 98.5 | 97.5 | 98.5 | 97.7 | 98.5 | 97.7 | 98.7 | 97.8 | 98.1 | 97.1 |
| 25 | 98.6 | 98.1 | 98.6 | 98.2 | 98.6 | 98.2 | 98.9 | 98.3 | 98.4 | 98.0 |
| 30 | 98.6 | 98.5 | 98.6 | 98.5 | 98.7 | 98.5 | 98.9 | 98.5 | 98.5 | 98.2 |
| 50 | 98.9 | 98.8 | 98.9 | 98.8 | 99.0 | 98.8 | 99.0 | 98.9 | 98.8 | 98.4 |
| 75 | 99.1 | 99.0 | 99.1 | 99.0 | 99.2 | 99.0 | 99.3 | 99.0 | 99.0 | 98.9 |
| 100 | 99.3 | 99.3 | 99.3 | 99.3 | 99.3 | 99.3 | 99.4 | 99.3 | 99.1 | 99.1 |
| 150 | 99.5 | 99.4 | 99.5 | 99.4 | 99.5 | 99.4 | 99.5 | 99.4 | 99.5 | 99.3 |
| 200 | 99.5 | 99.4 | 99.5 | 99.4 | 99.5 | 99.5 | 99.5 | 99.5 | 99.5 | 99.3 |

**Supplementary Table 4:**
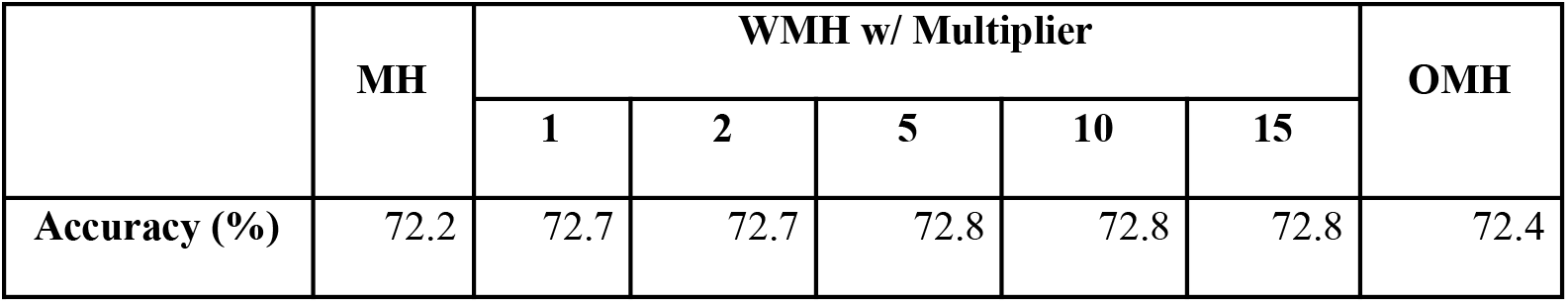
Effect of weight and order on MinHash accuracy. The inclusion of weight sees slight increases in performance over unweighted MinHash, while the addition of order results in smaller improvements.

**Supplementary Table 5:** Overhead of proposed sketching and sampling approaches. We count the number of 21-mers stored in the screen and the number of comparisons performed while classifying simulated 10Kb, 1% error, 10x coverage reads from the CGR microbial community, which has 4.85GB of sequence across 1,310 organisms. The screens are designed to capture 100 target matches per read. The sketching and sampling approaches only use a fraction of the unique k-mers present in the data, and thus have significantly reduced overhead compared to an exhaustive approach that stores all unique k-mers in the source genomes and looks up every k-mer present in the read sets. Note that the number of lookups performed in the exhaustive approach is an estimate, as the exhaustive approach is impractical to run in practice.

| Approach | Number of k-mers stored | # Lookups performed during Classification |
| --- | --- | --- |
| Exhaustive | 1,600,728,9970 | ~48,000,000,000 |
| MinHash | 60,382,132 | 604,985,465 |
| Minimizer | 69,382,660 | 687,003,545 |
| Uniform | 61,098,861 | 608,955023 |

**Supplementary Table 6:** Comparison of the uncompressed database sizes for the three index-based approaches. All three “Standard” databases are available directly from the developers of the tools. The pre-prepared Kraken2 and Centrifuge indexes cover all bacteria, archaea, viral and human genomes in Refseq, and the CLARK index is a complete set of bacterial genomes. The “custom” databases are further augmented to explicitly include the sequences in the CGR dataset, by adding the sequences and their taxonomy IDs to the existing database. This augmentation does not seem to add a significant amount of sequence.

| Approach | Size on Disk (GB) |
| --- | --- |
| Kraken2 (Standard) | 58.0 |
| Kraken2 (Custom) | 60.2 |
| Centrifuge (Standard) | 7.9 |
| Centrifuge (Custom) | 8.2 |
| CLARK (Standard) | 94.0 |

**Supplementary Table 7:** Time taken (computed as the “real” time from Unix’s “time” command) to classify simulated reads in our contaminant detection experiments, using 10Kb, 1% error, 10x coverage reads drawn from the CGR microbial dataset and the human reference genome GRCh38 (totalling ~7.85 Gb of sequence). This does not include the index-generation time for Kraken2 or Centrifuge, whose indexes were downloaded from the developers. All approaches are run with 32 threads. From these results, the speedup of index-based approaches over Minimap2 is clear, and as Minimap2 is already significantly faster than full sequence-to-sequence alignment, the limitations of alignment-based methods for large datasets should be evident. Within the two index-based approaches, Kraken2 is considerably faster, though it does require a significantly larger database (**Supplementary Table 6**).

| Approach | Runtime (minutes) |
| --- | --- |
| Minimap2 | 4680.3 |
| Kraken2 | 120.2 |
| Centrifuge | 410.7 |

